# How many genes can CRISPR edit to engineer complex adaptations?

**DOI:** 10.64898/2026.05.21.726991

**Authors:** Jinseul Kyung, Maliheh Esfahanian, Joseph Mann, Emily Koke, Keegan Pham, Yunru Peng, Moises Exposito-Alonso

## Abstract

Polygenic traits require the coordinated effects of multiple genes. Such complex traits have been a long-term target of study for geneticists, but multiplex CRISPR—the editing of multiple loci in the genome via multiple guide RNAs—is in its infancy. Reviewing 106 plant studies using multiplex CRISPR, we find that the multiplexing capacity has doubled every 5.4 years. Furthermore, a systematic experiment with 8, 16, and 24 simultaneous targets in *Arabidopsis thaliana* reveals efficiency of up to 75% in 24-plex editing in transformed plants confirmed by sequencing. We surprisingly found that the level of multiplexing, or the number of the targets, causes lesser efficiency reductions than other uncontrolled factors such as gRNA design or natural variation across plants. In fact, mathematical modeling of the decay in editing efficiency as a function of the gRNA numbers, showed a logistic model with sustained efficiency fits the data better than decay due to Cas9 competition or random editing. We then project that editing close to 100 genes in a plant can be feasible with reasonably large plant screens. However, feasible and reliable polygenic genome engineering will need developments outside of the CRISPR editing machinery itself, including innovations in gRNA vector delivery for large cargos, and a broader conceptual shift toward population-level poly-gene editing leveraging large distributions of mutations for breeding, natural selection, or experimental evolution.

**Author Contributions:** M.E.-A. conceived the project and secured funding. M.E.-A. and M.Es. designed the experimental strategy. M.Es. established the multiplex CRISPR and transformation pipelines in the laboratory, propagation through the T1 and T2 generations, and oversaw the first amplicon sequencing. Y.P. established the in-house iSeq amplicon sequencing protocol and contributed to cloning and genotyping pilots. M.Es supervised K.P. to construct cloning, bacterial transformations, plant growth, floral-dip transformations, and selection of T1 plants. E.K. contributed to early amplicon genotyping. J.K. propagated and sampled the T3 and J.K. and J.M. conducted the final amplicon sequencing panel. J.K. and M.E.-A. performed gene editing variant mapping, dataset quality control, and summarized results from published multiplex CRISPR studies. M.E.-A. modeled editing efficiency. J.K and M.E.-A. generated figures and wrote the first draft. All authors revised and improved the manuscript. J.K. and M.Es. contributed equally to this work and are designated as co-first authors.

## Introduction

Most traits of agronomic and evolutionary importance, such as yield, drought tolerance, flowering time, and disease resistance, are polygenic, shaped by the combined effects of tens to hundreds of loci across the genome (1). Engineering such traits therefore requires simultaneous modification of multiple genes, a challenge that single-gene knockouts cannot address. The ambition to edit polygenic architectures is not limited to crops: efforts to resurrect extinct species, such as the woolly mammoth by Colossal Biosciences, depend on introducing dozens of coordinated edits into a host genome (2). In plants, the BREEDIT project has begun targeting up to 60 growth-related genes in maize through multiplex CRISPR combined with doubled haploid technology (3). These examples underscore that the future of genome engineering is inherently multiplexed.

CRISPR is a genome editing system consisting of a guide RNA (gRNA), which directs the Cas9 protein to a specific DNA sequence through complementary base pairing, where Cas9 introduces a double-strand break. Target recognition additionally requires a short conserved sequence adjacent to the target site known as the protospacer adjacent motif (PAM), which is recognized directly by Cas and is essential for its binding and cleavage activity. The cell then repairs this break either by NHEJ, which disrupts gene function, or HDR, which enables precise sequence replacement. By simply redesigning the gRNA, virtually any PAM-proximal genomic locus can be targeted, making CRISPR a highly versatile tool for gene editing.

CRISPR-Cas9, since its first application in plants in 2013 (4), has rapidly become the dominant tool for targeted mutagenesis. An immediate extension was to deliver multiple single guide RNAs (sgRNAs) in tandem arrays, enabling simultaneous editing of several loci in a single transformation event. Technical innovations have steadily expanded the toolkit: polycistronic tRNA-gRNA (PTG) systems streamlined sgRNA expression (5), ribozyme-based strategies improved editing precision (6), engineered Cas9 variants increased specificity (7), and Cpf1/Cas12a added self-processing capabilities (8). Delivery methods, including Agrobacterium-mediated transformation, protoplast transfection, and nanoparticle-based approaches, have improved across diverse species (9), while base-editing and prime-editing systems have extended the precision of modifications beyond simple knockouts (10). Computational design tools have further helped minimize off-target effects (11).

Yet the realities of polygenic architectures demand multiplexing at a scale that current practice has barely approached. If hundreds of loci contribute to a trait of interest, can multiplex CRISPR realistically target them all? Most published experiments still use fewer than five sgRNAs, and the relationship between the number of targets and per-site editing efficiency remains poorly characterized. It is unclear whether efficiency decays linearly with additional targets — as expected from simple sgRNA competition for a limited Cas9 pool — or whether more complex dynamics, such as synergistic chromatin opening, in which simultaneous targeting of nearby genomic regions by multiple gRNAs collectively loosens local chromatin structure and enhances accessibility beyond what a single guide could achieve, might buffer the decline.

Here we address this question through a meta-analysis of 106 published multiplex CRISPR studies in plants (123 independent experiments) and a systematic experimental evaluation of editing efficiency at 8, 16, and 24 simultaneous target sites in *Arabidopsis thaliana*. We find that the frontier of multiplex editing has grown exponentially over the past decade, with a doubling time of approximately 5 years. Experimentally, targeting multiple genes, or high-level multiplexing, is achievable: up to 73% of plants that targeted 24 sites spread across eight genes, carried mutations in all eight targeted genes. We used binomial generalized linear mixed models to quantify the sources of variation in editing success and to predict efficiency at higher target numbers. Our analysis reveals that gRNA identity but not multiplex level is the dominant source of variation, and that the observed efficiency decay is better described by a diminishing-interference model than by linear efficiency decay or competition for Cas9. Together, these results provide a quantitative framework for the design and interpretation of high-level multiplex CRISPR experiments in plants and other organisms.

### CRISPR multiplex for plants is on the rise

To evaluate the current landscape of multiplex CRISPR use in plants, we reviewed 106 papers encompassing 123 independent experiments (**Table S1**). Literature was retrieved using the keywords “multiplex,” “CRISPR,” “plant”, and article type= “Research Support” in Pubmed curated manually. Since the foundational study by Li et al (4), the number of sgRNAs used per experiment has increased substantially over the past decade (**Fig. 1**).

**Figure 1.**
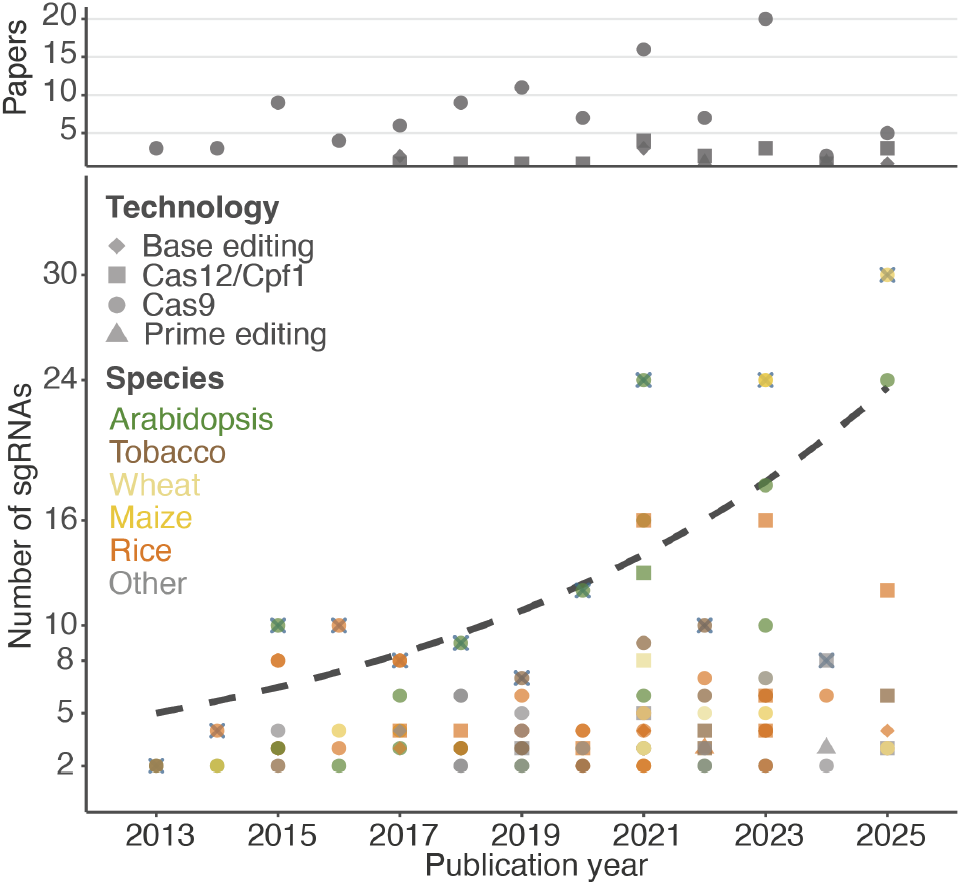
Review of multiplex CRISPR shows exponential use over time. (Upper) Number of papers reviewed containing multiplex CRISPR approaches split by technology used. (Lower) Year of publication and number of sgRNAs used across plant species. The dashed line represents an exponential model of the maximum number of gRNAs used per year, corresponding to a doubling time of 5.4 years. (file: illustrator/Fig1.png, half page)

While early studies typically used just two sgRNAs (2× multiplexing), more recent work has reached 24× in *Arabidopsis thaliana* and 30× in maize. Since 2014, maize has emerged as a frequent platform for high-level multiplexing due to strong demand for complex trait engineering in this widely cultivated crop. Nevertheless, the majority of studies (88 out of 123) to date still cluster around 4× multiplexing, suggesting that high-degree multiplexing remains technically challenging or underutilized.

We also observed diversification in the target species over time. Early efforts focused primarily on model species, especially *Arabidopsis*. However, the number of targeted species has increased gradually. Correspondingly, multiplex CRISPR has been applied to major crops including maize and wheat. Among these, rice stands out, appearing in 23 independent studies—highlighting its experimental tractability and agricultural importance. In the 2020s, the technique has extended to additional species such as cotton and tomato, indicating broader potential for agricultural applications. Examining papers for the number of transformed plants, selection-marker-filtered lines, and sequenced lines, we could not generate standardized efficiency metrics (**Table S1**). While this likely reflects that current literature is mostly focused on generating useful CRISPR mutations for biological studies rather than studying efficiency per se, we call for more standardized efficiency reporting of constructs.

In summary, plant multiplex CRISPR research has advanced steadily in both technical complexity and species diversity. However, high-level multiplexing (beyond 4 sgRNAs) is still relatively rare, pointing to the need for systematic evaluation of its efficiency and limitations.

### High editing efficiency is maintained even up to 24 targets

To better address how editing efficiency scales with the number of genomic targets, we conducted our own systematic experiment. We evaluated multiplex CRISPR systems targeting 8, 16, or 24 genomic sites across 8 genes (*DOG1, EAR1, FLC, GL1, IND, PAN, PCO5, PAN, PCO5, TT8*). Target genes are selected for future phenotypic analysis, but no phenotypic selection is performed here. We employed a previously published system by Stuttmann et al. (12) (Addgene pDGE construct: https://www.addgene.org/browse/article/28211370/) that delivers all sgRNAs through a single construct containing a tandem sgRNA array driven by universal Pol III promoters and a short poly-T stretch of terminators. CRISPR constructs were introduced into Col-0 plants via *Agrobacterium*-mediated transformation, and approximately one hundred progeny were propagated over three generations. Gene editing outcomes were validated using Illumina amplicon sequencing and mutation profiling with CRISPResso (see **Methods, Table S2**). This gave us a rich database of total 4224 genomic regions (24 loci in 176 sequenced plants) to study PAM-site mutations and drivers of efficiency (**Fig. 2**).

**Figure 2.**
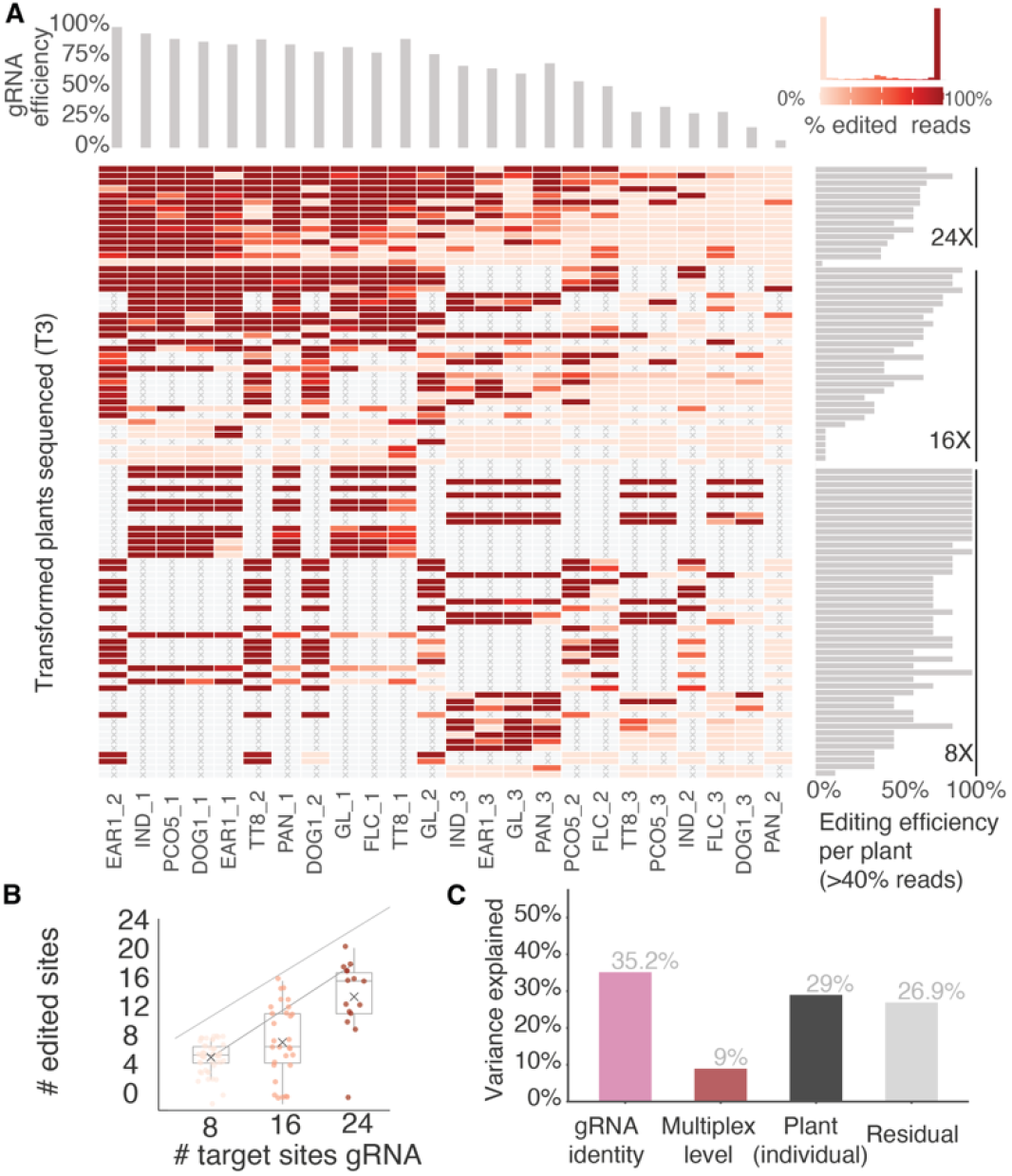
Multiplex editing efficiency is high even at 24X. (A) Heatmap indicating the percentage of short read sequencing support for a mutation nearby the PAM site target for each plant and each of the up to 24 target sites. Marginal histograms indicate the efficiency of 24 gRNA across all plants and efficiency of edits in each of the 92 T3 plants across all sites. A site is considered edited if 30% of the reads support the non-reference allele, a threshold that was defined empirically (see inset of percentage of edited reads). (B) The total number of edited sites across the three levels of gRNA multiplex. (C) Sources of variance in editing efficiency explained in a Generalized Linear Mixed Model by different factors. (file: illustrator/Fig2.png, half page)

We first examined overall mutation rates using coverage of reads mapping to the amplicon reference region and called it as an alternative allele by CRISPResso (**Fig. 2A, Fig. S1**). With abundant sequencing coverage on the amplicon (mean=10,285 average fold depth), we were able to detect that most loci had either reference or alternative homozygote genotypes, with a few loci showing heterozygote frequencies (**Fig. 2**), likely owed to the continued Cas9 activity (transformed constructs were deliberately not segregated out and negatively selected). With this, we found that with increasing multiplex we expectedly increased the number of target sites edited, with a slope = 0.413 ± 0.062 sites per added target (*P* = 1.8×10^-9^, *R*^*2*^ =0.33) (**Fig. 2B**)

We first assessed whether there was a mutation overlapping the PAM site, which were edited at a mean efficiency of 57.6% ± 26.5% across guide RNAs (n = 24) and 61.2% ± 27.1% across 110 sequenced plants. For the highest level multiplex, 24X, even with only 12 independent transformants and no pre-selection beyond construct presence with Basta, we found a T3 plant with 19 independent PAM site edits.

Although the overall efficiency is more dependent on the nature of the PAM sequence rather than the nature of the specific gene, many researchers use multiplex CRISPR with redundant gRNAs to increase the likelihood that the desired gene is edited. To test the impact of multiple guides per gene to create KOs, we designed 8X, 16X, and 24X constructs to target the same 8 genes. With a single gRNA per gene (8X), we achieve a complete 8 KO (octuple mutant) set on 21% of plants. With two gRNAs per gene (16X) we achieved octuple mutants in 37% plants, and with three gRNAs (24X) we achieved 73%. These results support the common rule of thumb in the literature that having multiple gRNAs per gene increases the probability of generating knockout alleles.

Finally, we wanted to study how much of the variation in efficiency may be attributed to different factors, such as specific gRNAs, specific transformed plants, or multiplexing level. A Generalized Linear Mixed Model (GLMER) with presence/absence as a response variable attributed the largest share of variation to gRNA identity, in other words, PAM sequence itself (35.2%), followed by plant-to-plant differences (24.4%), with the multiplex level itself accounting for 8.0% of variation (**Fig. 2C**). This emphasizes that guide sequence quality is often more limiting than the high-level multiplexing,

### Towards a hundred genes: potential efficiency under very-high-multiplex CRISPR

Keeping in mind that gRNA design and uncontrolled plant biological factors explain much of the observed editing efficiency variation, a key question is whether CRISPR multiplexing will be able to alter polygenic traits considering an efficiency decay with larger numbers of genes or loci. While we intuitively expect that the gene editing efficiency would decrease as we increase the number of the targets and this is often discussed among colleagues attempting large mutation arrays, we are not aware of principled models of efficiency loss in CRISPR multiplex literature. To formalize the relationship, we compared three simple theoretical null models and three empirically fitted decay models, and tested their explanatory power of the data (**Fig. 3A**). The three analytical null expectations include: (**M1**) **no decay**, where average editing efficiency of one gRNA is maintained no matter how many other gRNAs are co-transformed. We used as possible ranges *p*_*edit*_*=*70%–80% efficiency reported in the literature and similar to our gRNA average 73% efficiency (pseudo-R^2^ =−0.27, AIC = 985.2, ΔAIC = 439.3). (**M2**) **joint independent probability** model simply considers the fact that two events of a given probability (e.g. *p=*70%) to happen jointly multiply for as many *n* gRNAs as there are in the construct: *P(site* |*n) = p* ^*n*^; which expects a fast efficiency decay and does not fit the data well (pseudo-R^2^ = 0.04, AIC = 753.5, ΔAIC = 207.6). (**M3**) **Cas9 competition model**, which captures a popular belief that editing efficiency may be limited by Cas9 expression of the plant. This model still explained the data poorly (pseudo-R^2^ = −0.09, AIC = 852.5, ΔAIC = 306.6). We then consider four alternative models that are fitted to the data. (**M4**) **Naïve logistic fit** to empirical dataset following *P(site* |*n) = 1 · logistic(a + b n)* (pseudo-R^2^ = 0.04, AIC = 752.9, ΔAIC = 207.0). Because this model assumes efficiency decays constantly with increasing *n* number of gRNAs, it quickly predicts efficiency should plateau at zero. Because we observe a decay in efficiency but not as steep as expected for 24X, we proposed (**M5**) a **logistic fit with diminishing decay** that expects efficiency decays following *log(n)*, which improved predictability (pseudo-R^2^ = 0.06, AIC = 738.8, ΔAIC = 192.9). This model still assumes efficiency would eventually approximate zero, thus we implemented a (**M6**) **logistic diminishing decay with minimum efficiency**, which enforces a floor of efficiency of at least 1% (pseudo-R^2^ = 0.06, AIC = 738.7, ΔAIC = 192.8). The floor percentage could be attempted to be modeled as a hyperparameter, but for this to be accurate we would need empirical multiplex data at 100x gRNAs, which was not possible. The model was not generally sensitive to low percentage digits (e.g. 1% or 2% not shown), as long as it is non-0% which is the reason we chose this over M4-5 that assumes 0%, which is not biologically realistic. Finally, because of the broad efficiency differences observed across plants, we implemented the (**M7**) **logistic decay with minimum efficiency and overdispersion**, to capture the broader observed variance (pseudo-R^2^ = 0.31, AIC = 545.9, ΔAIC = 0.0), the best-supported model by a wide margin. There may be multiple mechanisms involved in this phenomenon, but we speculate this diminishing decay model is consistent with a hypothesis of Cas9 activity at early targets opens chromatin or activates DNA damage response pathways, partially offsetting the cost of additional targets (13).

**Figure 3.**
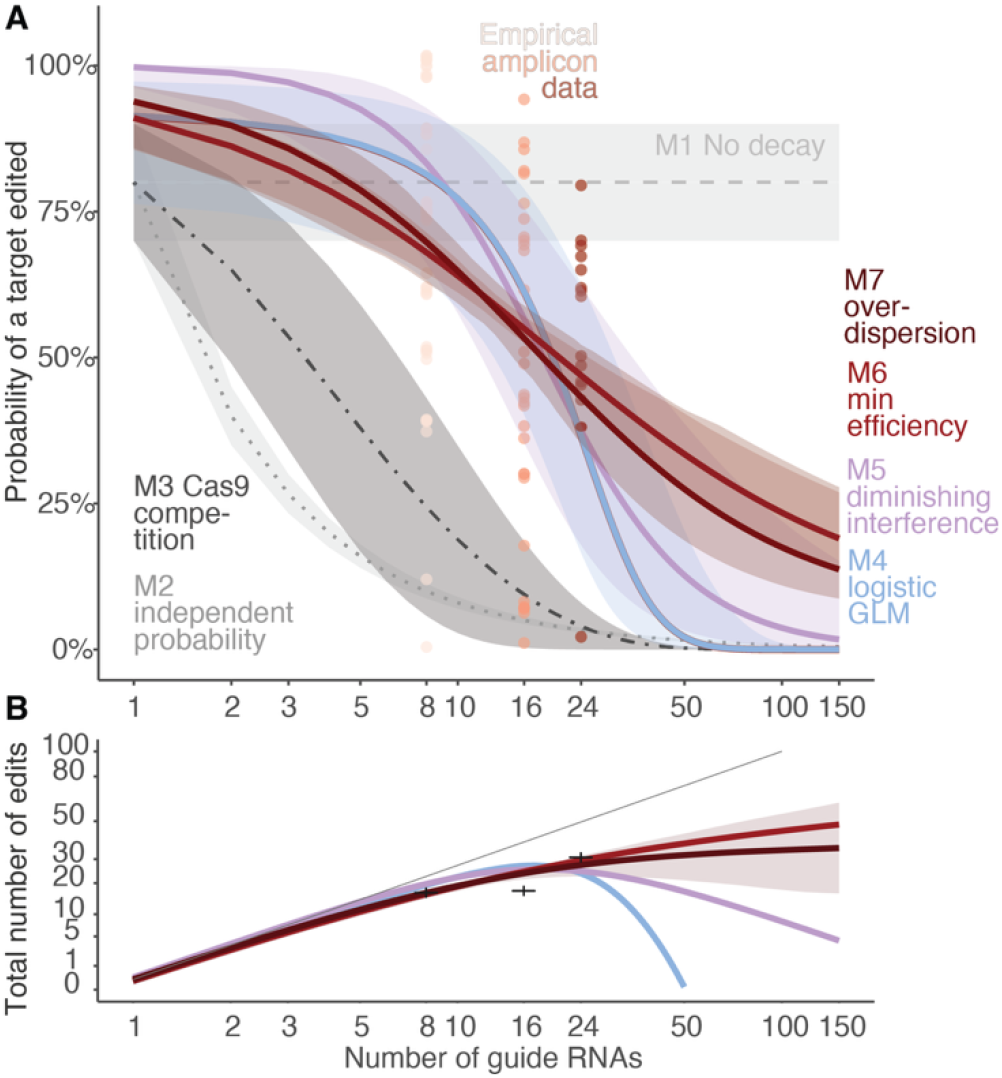
Models of multiplex CRISPR efficiency and projections. (A) Description of possible models of efficiency decay of multiplex-CRISPR with increasing gRNA and genome targets. Horizontal line indicates no decay model (M1), dotted line and light grey confidence interval indicates independent joint probability (M2), dark gray indicates Cas9-like competition (M3). (B) Total number of targets edited with increasing number of gRNAs. Most complex models are displays (M4-6) as well as no-decay null model for comparison. (file: illustrator/Fig3.png, half page)

Then we play the *gedanken “*thought” experiment: how feasible would it be to edit 100 genes at once? Since polygenic architectures are probably comprised by several hundred coregenes (1, 14, 15), we can use the best fit efficiency model (M7) to predict *P(site* | *100) = 0*.*01 + 0*.*99 · logistic(2*.*86 – 0*.*95 · log 100) =* 18.7% average efficiency for any of the 100 gRNA targets and an average plant would contain only 18 hits, but the top 99.99% plant—i.e. the top plant out of 10,000 offspring plants, assuming 10 antibiotics selected seeds per 1,000 transformed plants—would contain ∼94 genome target edits. While a large and intensive experiment, in principle, this would be achievable.

### Outlook and perspectives

Have these experiments then solved the challenge of polygenic gene editing? Not quite. It is likely that new technologies will need to be developed for polygenic editing to be routinely conducted in plant molecular genomics. For instance, the plant expression vector system used here (12) can accommodate approximately 40–50 gRNAs within the T-DNA size limit of standard binary vectors (∼25 kb) for *Agrobacterium-*mediated transformations.

Innovations in larger-capacity vectors or optimizing (16), co-transformation of multiple *Agrobacterium* strains (17), or alternative delivery systems may be needed. For instance, a smaller editor than Cas9 (18) may provide further space for cargo, but this may be limited. A viral vector could deliver 6 gRNAs without the need for multi-generational transformations if multiple viral particles infect stem cells (19, 20). Automation of detection via amplicon sequencing of many plants x amplicon combinations would be a further aid in experimental pipelines.

Ultimately, the community may need to re-consider how molecular biology studies the consequences of gene edits and mutations. By definition, polygenic architectures will be controlled by many genes which may communally contribute to phenotypic effects of a quantitative or continuous trait, and none of the edits may be the holy grail genome change. Therefore, multiplex CRISPR experiments may seek, instead of a full complement of 100 mutations, a population of plants with different subsets of targeted mutations that would display a distribution of a quantitative trait, which can then be selected by classic breeding (3) or experimental evolution (21).

Two decades after CRISPR entered plant biology, the question is no longer *whether* multiplex editing works, but *how far it can scale* — and what kind of science it enables once it does. Our review analysis shows the field doubling its multiplex frontier every five years; our 24× experiments show that complete octuple knockouts are already routine; and our best-supported model predicts that a realistic plant screen could deliver a single individual edited at ∼95 of 100 target genes. Crossing the gap to a hundred clean knockouts is an engineering problem, but it is no longer a conceptual one. Multiplex CRISPR will not only scale genome engineering; it will reshape how we interrogate the polygenic architecture of life itself.

## Methods

### Generation of Multiplex Mutants

Eight *Arabidopsis thaliana* genes— *DELAY OF GERMINATION 1* (DOG1; AT5G45830), *ENHANCER OF ABA CO-RECEPTOR* 1 (EAR1; AT5G22090), *FLOWERING LOCUS C* (FLC; AT5G10140), *GLABRA 1* (GL1; AT3G27920), *INDEHISCENT* (IND; AT4G00120), *PERIANTHIA* (PAN; AT1G68640), *PLANT CYSTEINE OXIDASE 5* (PCO5; AT3G58670), and *TRANSPARENT TESTA 8* (TT8; AT4G09820)—were selected for multiplex CRISPR targeting. For each gene, three sgRNAs were designed using the CHOPCHOP web tool (https://chopchop.cbu.uib.no) (22).

Multiplex CRISPR constructs were generated following the protocol of Stuttmann et al. (12). Briefly, three separate transcriptional arrays of eight sgRNAs were individually assembled into intermediate vectors, producing three 8× constructs. These transcriptional units were then assembled into final binary vectors using GoldenGate cloning to generate 16× and 24× constructs. Final vectors were introduced into *A. thaliana* via the Agrobacterium mediated floral dip method. Transgenic plants were selected by resistance to Basta, and T2 and T3 generations were used for downstream analysis.

### Plant transformation

Binary vectors were initially cloned in *E. coli* and then transformed into *Agrobacterium tumefaciens strain GV3101* for stable plant transformation via floral dip. Through the antibiotics selection, we selected 12 randomly selected T1 plants per each construct, without any phenotypic selection. And propagated into T2 plants. T3s were derived from T2s and sometimes we did quality control checks sequencing both T2 and T3 of the same transformed T1 lineage (Fig. S2). For several T2s we only sequenced until T2 since T3 plants were unavailable.

### Amplicon sequencing

For each construct, 12 T2 and 8-18 T3 transgenic plants were randomly selected. For T3 plants, the number of the plants differed due to the loss during the germination or survival. Genomic DNA was extracted using a modified CTAB protocol (23, 24).

Genotyping was performed via two-step amplicon PCR (25). In brief, in the first PCR, gene-specific primers flanking the sgRNA target regions were used along with partial adapter sequences. In the second PCR, full-length Illumina Nextera P5/P7 adapters and unique dual barcodes were incorporated. Indexed libraries were pooled and sequenced on an Illumina iSeq 100 platform.

Amplicon sequencing was performed across four sequencing rounds over the course of the project. The first round (312 samples) established the amplicon-PCR and iSeq workflow on T1 and T2 plants. A second round (312 samples) was subsequently re-sequenced with corrected library indices in a third round (identical 312 samples) to obtain a usable dataset for these plants. The fourth and final round, dataset reported in this manuscript was generated on the T3 generation: 92 plants were each amplified at 16 target regions across 8 genes, yielding 1,472 indexed amplicon libraries pooled across four iSeq 100 runs (Q30 ≥ 85% per run, with even read distribution across targets). Lineage and per-generation sequencing counts are summarized in Table S2.

### Mutation calling and Efficiency calculation

Sequencing data were analyzed using CRISPResso2 (26), integrated with a custom Snakemake pipeline and Python scripts for automated processing. Read alignment, mutation calling, and coverage statistics were extracted from CRISPResso2 output files. A target site was called as mutated if ≥30% of sequencing reads showed modification (indels or substitutions) relative to the reference sequence at the gRNA cut site.

### Statistical modeling of editing efficiency

To model how per-site editing probability changes with the number of target sites, we compared five models fit to 1,216 individual plant × targeted site observations from T3-generation plants. The response variable was binary (mutated or not at ≥30% modified reads). All GLMMs were fit using lme4::glmer() in R with Laplace approximation. Models 1–3 are analytical null expectations evaluated at two baseline per-gRNA efficiencies: *p*_0_ = 0. 70 (literature median) (27, 28). and *p*_0_ = 0. 90 (GLMM extrapolation to *n* = 1), shown as bands between the two baselines.

#### Model 1 — No decay (constant efficiency)

Per-site editing probability is independent of the number of targets:

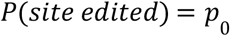

#### Model 2 — Cas9 competition (1/n decay)

A fixed pool of Cas9 protein is shared equally among all *n* target sites:

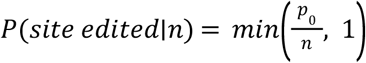

#### Model 3 — Independent joint probability (*p*^*n*^)

Each of *n* sites is edited independently with probability *p* ; the probability that *all* sites are simultaneously edited is:

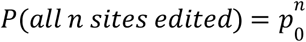

This joint probability is shown for reference to illustrate the combinatorial challenge of complete editing, even without per-site decay.

#### Model 4 — Binomial GLMM with linear efficiency decay

A generalized linear mixed model with binomial family and logit link:

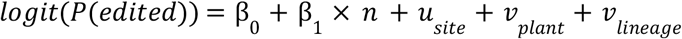

where *n* is the total number of target sites,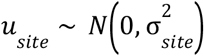 captures gRNA-specific variation, and 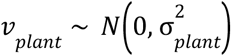 captures plant-to-plant variation. This model assumes each additional gRNA imposes the same constant penalty on the log-odds scale. 95% confidence intervals were computed via the delta method:

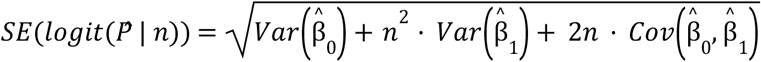

where 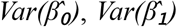 and 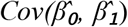 are taken from the estimated variance–covariance matrix of the fixed effects (vcov() of the fitted GLMM) and therefore already incorporate the sample size; no further division by 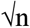 is required. The 95% CI is computed on the logit scale as 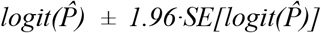 and back-transformed via the inverse logit.

#### Model 5 — Diminishing interference (log-decay)

The linear term of M4 in *n* is replaced by a logarithmic term:

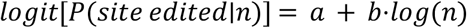

Each *doubling* of targets has the same effect on the log-odds of editing, regardless of whether going from 2 → 4 or 50 → 100. Biologically, this is consistent with early Cas9 activity opening chromatin or activating DNA-damage-response pathways that partially offset the cost of additional targets, analogous to facilitated recombination in DSB repair.

#### Model 6 — Log-decay with efficiency floor

Because M5 implies efficiency approaches zero at very high *n*, which is inconsistent with the long-standing floor of residual editing observed in saturation experiments, we added a fixed minimum efficiency *c*:

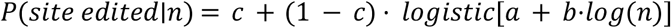

with *c* = 0.01. This enforces a biologically plausible asymptote of 1% efficiency without altering behavior at low to moderate *n*.

#### Model 7 — Beta-binomial with log-decay and floor (overdispersion)

Substantial plant-to-plant variation in editing competence remained unexplained by the binomial models M4–M6 (residual deviance > d.f.). We therefore replaced the Binomial likelihood with a Beta-Binomial, allowing each plant’s per-site probability to be drawn from a Beta distribution with mean *μ(n)* and intraclass correlation *ρ*:

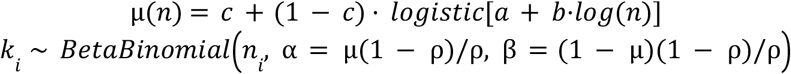

where *k*_*i*_ is the number of edited sites out of *n*_*i*_ in plant *i*, and *c = 0*.*01* as in M6. The overdispersion parameter *ρ* captures the extent to which plants differ in their overall editing competence beyond what a single shared probability predicts. M7 improved fit by **ΔAIC ≈ 193** over M6 (see Table S3).

#### Projections to higher multiplex levels

Per-site probabilities at *n* = 100 or 300 were obtained by substituting *n* directly into the fitted M7 mean function. Distributions of edited sites (or genes, when multiple gRNAs per gene were modeled) across h *f* T3ypothetical 10,000-plant T3 screens were obtained by Monte Carlo simulation: for each plant, a per-site probability was drawn from the Beta distribution implied by *μ(n)* and *ρ*, then site-level outcomes were drawn Binomially, and (for multi-gRNA-per-gene designs) a gene was counted as edited if at least one of its gRNAs hit. Quantiles and maximum order statistics reported in the Discussion are empirical summaries of 10^6^-plant simulations.

### Variance decomposition

A separate Binomial GLMM was fit with random intercepts for multiplex level, gRNA identity, and plant individual (no fixed effect for multiplex level):

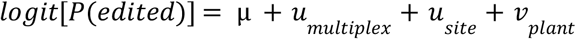

Variance components were extracted on the logit scale. The residual (level-1) variance for a Binomial GLMM with logit link is fixed at *π*^*2*^*/3* ≈ 3.29 (the variance of the standard logistic distribution). The percentage of variance explained by each component is:

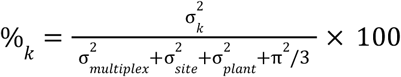

### Apparent editing at non-targeted sites

In constructs with fewer than 24 gRNAs (8× and 16×), some non-targeted sites within the same gene showed modification above the calling threshold. All 86 such events occurred at sites within the same gene as a targeted site—never at a completely untargeted gene. Two mechanisms account for this: (1) *Amplicon overlap*: For most genes (GL, FLC, PCO5, IND, PAN, EAR1, DOG1), the three gRNA target sites per gene are located 3–60 bp apart on the same or overlapping PCR amplicons. A typical Cas9-induced deletion of 10–25 bp at the targeted cut site physically spans the neighboring non-targeted gRNA sequence. CRISPResso2 scores both sites as modified from the same sequencing reads, representing a single deletion event detected at two nearby positions rather than independent editing. (2) *PCR amplicon dropout (TT8)*: The three TT8 target sites are located 491–3,743 bp apart on separate PCR amplicons. In some 8× plants, the non-targeted TT8 amplicons showed very low read counts (>99.5% of reads fail to align) but high apparent modification percentages. This is consistent with a large deletion at the targeted site extending into the primer binding region of a neighboring amplicon, preventing PCR amplification. These observations do not affect the GLMM projections, which are fit exclusively to targeted site data. However, they highlight that amplicon-based genotyping can overestimate editing at closely spaced sites.

## Additional information

## Acknowledgements

We thank Benjamin Jin for their contributions to plant care and laboratory support during the project periods, and members of the MOILAB for discussion and manuscript review: Ruth Epstein, Meixi Lin, Miles Roberts, Benjamin Jin. **Funding** M.E.-A. is supported by the Office of the Director of the National Institutes of Health’s Early Investigator Award (1DP5OD029506-01), the U.S. Department of Energy, Office of Biological and Environmental Research (DE-SC0021286), by the U.S. National Science Foundation’s DBI Biology Integration Institute WALII (Water and Life Interface Institute, 2213983), by the Carnegie Institution for Science, the Howard Hughes Medical Institute, the Innovative Genomics Institute, and the University of California Berkeley. Computational analyses were done on the High-Performance Computing clusters of the Carnegie Institution for Science and High Performance Computing cluster of the University of California Berkeley.

## Data availability

All Illumina sequencing reads are available at NCBI with project: XXX, all scripts are available at Zenodo: <DOI add>.

## Supplemental Materials

**Table S1.**
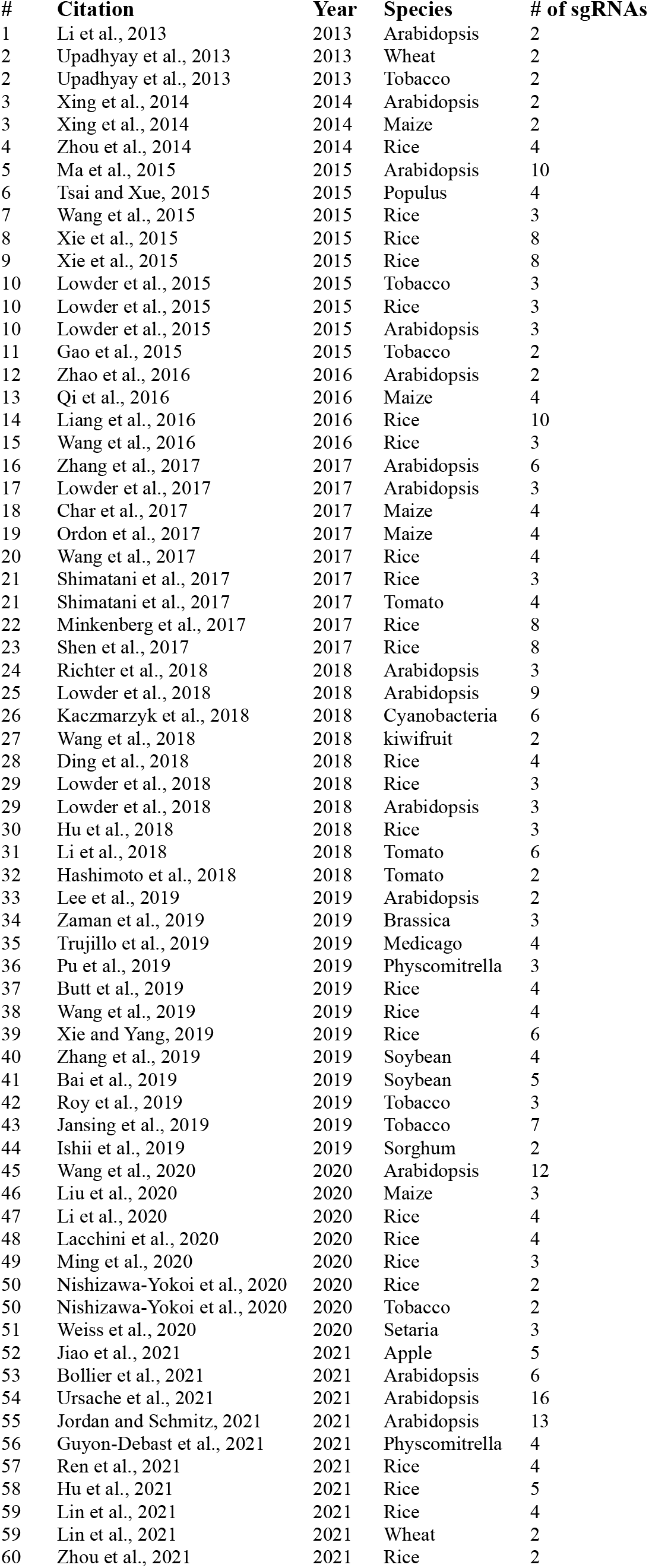
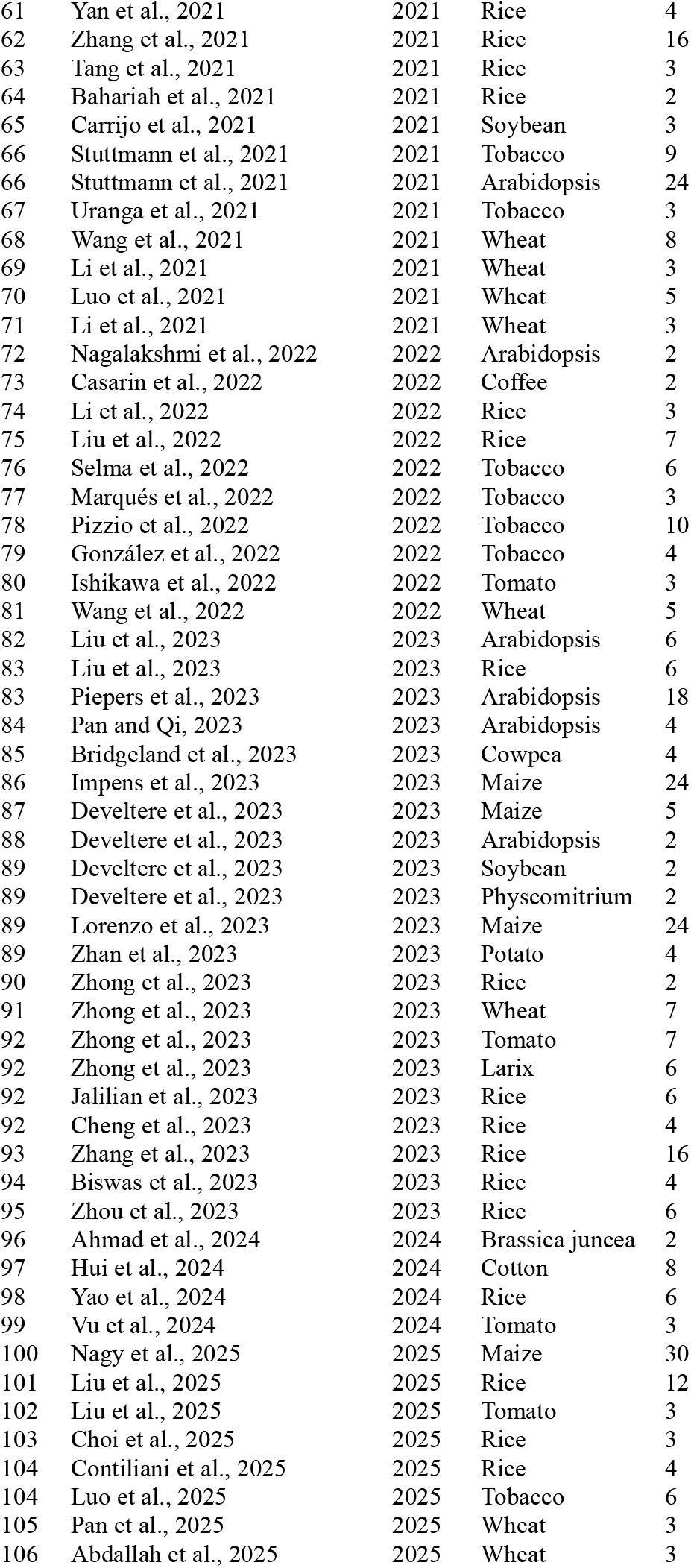
Meta-analysis of multiplex CRISPR studies in plants. Summary of published experiments using multiplex CRISPR methods in plants. Each entry includes the plant species, number of sgRNAs used, and publication year.

| # | Citation | Year | Species | # of sgRNAs |
| --- | --- | --- | --- | --- |
| 1 | Li et al., 2013 | 2013 | Arabidopsis | 2 |
| 2 | Upadhyay et al., 2013 | 2013 | Wheat | 2 |
| 2 | Upadhyay et al., 2013 | 2013 | Tobacco | 2 |
| 3 | Xing et al., 2014 | 2014 | Arabidopsis | 2 |
| 3 | Xing et al., 2014 | 2014 | Maize | 2 |
| 4 | Zhou et al., 2014 | 2014 | Rice | 4 |
| 5 | Ma et al., 2015 | 2015 | Arabidopsis | 10 |
| 6 | Tsai and Xue, 2015 | 2015 | Populus | 4 |
| 7 | Wang et al., 2015 | 2015 | Rice | 3 |
| 8 | Xie et al., 2015 | 2015 | Rice | 8 |
| 9 | Xie et al., 2015 | 2015 | Rice | 8 |
| 10 | Lowder et al., 2015 | 2015 | Tobacco | 3 |
| 10 | Lowder et al., 2015 | 2015 | Rice | 3 |
| 10 | Lowder et al., 2015 | 2015 | Arabidopsis | 3 |
| 11 | Gao et al., 2015 | 2015 | Tobacco | 2 |
| 12 | Zhao et al., 2016 | 2016 | Arabidopsis | 2 |
| 13 | Qi et al., 2016 | 2016 | Maize | 4 |
| 14 | Liang et al., 2016 | 2016 | Rice | 10 |
| 15 | Wang et al., 2016 | 2016 | Rice | 3 |
| 16 | Zhang et al., 2017 | 2017 | Arabidopsis | 6 |
| 17 | Lowder et al., 2017 | 2017 | Arabidopsis | 3 |
| 18 | Char et al., 2017 | 2017 | Maize | 4 |
| 19 | Ordon et al., 2017 | 2017 | Maize | 4 |
| 20 | Wang et al., 2017 | 2017 | Rice | 4 |
| 21 | Shimatani et al., 2017 | 2017 | Rice | 3 |
| 21 | Shimatani et al., 2017 | 2017 | Tomato | 4 |
| 22 | Minkenberg et al., 2017 | 2017 | Rice | 8 |
| 23 | Shen et al., 2017 | 2017 | Rice | 8 |
| 24 | Richter et al., 2018 | 2018 | Arabidopsis | 3 |
| 25 | Lowder et al., 2018 | 2018 | Arabidopsis | 9 |
| 26 | Kaczmarzyk et al., 2018 | 2018 | Cyanobacteria | 6 |
| 27 | Wang et al., 2018 | 2018 | kiwifruit | 2 |
| 28 | Ding et al., 2018 | 2018 | Rice | 4 |
| 29 | Lowder et al., 2018 | 2018 | Rice | 3 |
| 29 | Lowder et al., 2018 | 2018 | Arabidopsis | 3 |
| 30 | Hu et al., 2018 | 2018 | Rice | 3 |
| 31 | Li et al., 2018 | 2018 | Tomato | 6 |
| 32 | Hashimoto et al., 2018 | 2018 | Tomato | 2 |
| 33 | Lee et al., 2019 | 2019 | Arabidopsis | 2 |
| 34 | Zaman et al., 2019 | 2019 | Brassica | 3 |
| 35 | Trujillo et al., 2019 | 2019 | Medicago | 4 |
| 36 | Pu et al., 2019 | 2019 | Physcomitrella | 3 |
| 37 | Butt et al., 2019 | 2019 | Rice | 4 |
| 38 | Wang et al., 2019 | 2019 | Rice | 4 |
| 39 | Xie and Yang, 2019 | 2019 | Rice | 6 |
| 40 | Zhang et al., 2019 | 2019 | Soybean | 4 |
| 41 | Bai et al., 2019 | 2019 | Soybean | 5 |
| 42 | Roy et al., 2019 | 2019 | Tobacco | 3 |
| 43 | Jansing et al., 2019 | 2019 | Tobacco | 7 |
| 44 | Ishii et al., 2019 | 2019 | Sorghum | 2 |
| 45 | Wang et al., 2020 | 2020 | Arabidopsis | 12 |
| 46 | Liu et al., 2020 | 2020 | Maize | 3 |
| 47 | Li et al., 2020 | 2020 | Rice | 4 |
| 48 | Lacchini et al., 2020 | 2020 | Rice | 4 |
| 49 | Ming et al., 2020 | 2020 | Rice | 3 |
| 50 | Nishizawa-Yokoi et al., 2020 | 2020 | Rice | 2 |
| 50 | Nishizawa-Yokoi et al., 2020 | 2020 | Tobacco | 2 |
| 51 | Weiss et al., 2020 | 2020 | Setaria | 3 |
| 52 | Jiao et al., 2021 | 2021 | Apple | 5 |
| 53 | Bollier et al., 2021 | 2021 | Arabidopsis | 6 |
| 54 | Ursache et al., 2021 | 2021 | Arabidopsis | 16 |
| 55 | Jordan and Schmitz, 2021 | 2021 | Arabidopsis | 13 |
| 56 | Guyon-Debast et al., 2021 | 2021 | Physcomitrella | 4 |
| 57 | Ren et al., 2021 | 2021 | Rice | 4 |
| 58 | Hu et al., 2021 | 2021 | Rice | 5 |
| 59 | Lin et al., 2021 | 2021 | Rice | 4 |
| 59 | Lin et al., 2021 | 2021 | Wheat | 2 |
| 60 | Zhou et al., 2021 | 2021 | Rice | 2 |
| 61 | Yan et al., 2021 | 2021 | Rice | 4 |
| 62 | Zhang et al., 2021 | 2021 | Rice | 16 |
| 63 | Tang et al., 2021 | 2021 | Rice | 3 |
| 64 | Bahariah et al., 2021 | 2021 | Rice | 2 |
| 65 | Carrijo et al., 2021 | 2021 | Soybean | 3 |
| 66 | Stuttman et al., 2021 | 2021 | Tobacco | 9 |
| 66 | Stuttman et al., 2021 | 2021 | Arabidopsis | 24 |
| 67 | Uranga et al., 2021 | 2021 | Tobacco | 3 |
| 68 | Wang et al., 2021 | 2021 | Wheat | 8 |
| 69 | Li et al., 2021 | 2021 | Wheat | 3 |
| 70 | Luo et al., 2021 | 2021 | Wheat | 5 |
| 71 | Li et al., 2021 | 2021 | Wheat | 3 |
| 72 | Nagalakshmi et al., 2022 | 2022 | Arabidopsis | 2 |
| 73 | Casarin et al., 2022 | 2022 | Coffee | 2 |
| 74 | Li et al., 2022 | 2022 | Rice | 3 |
| 75 | Liu et al., 2022 | 2022 | Rice | 7 |
| 76 | Selma et al., 2022 | 2022 | Tobacco | 6 |
| 77 | Marqués et al., 2022 | 2022 | Tobacco | 3 |
| 78 | Pizzio et al., 2022 | 2022 | Tobacco | 10 |
| 79 | González et al., 2022 | 2022 | Tobacco | 4 |
| 80 | Ishikawa et al., 2022 | 2022 | Tomato | 3 |
| 81 | Wang et al., 2022 | 2022 | Wheat | 5 |
| 82 | Liu et al., 2023 | 2023 | Arabidopsis | 6 |
| 83 | Liu et al., 2023 | 2023 | Rice | 6 |
| 83 | Piepers et al., 2023 | 2023 | Arabidopsis | 18 |
| 84 | Pan and Qi, 2023 | 2023 | Arabidopsis | 4 |
| 85 | Bridgeland et al., 2023 | 2023 | Cowpea | 4 |
| 86 | Impens et al., 2023 | 2023 | Maize | 24 |
| 87 | Develtere et al., 2023 | 2023 | Maize | 5 |
| 88 | Develtere et al., 2023 | 2023 | Arabidopsis | 2 |
| 89 | Develtere et al., 2023 | 2023 | Soybean | 2 |
| 89 | Develtere et al., 2023 | 2023 | Physcomitrium | 2 |
| 89 | Lorenzo et al., 2023 | 2023 | Maize | 24 |
| 89 | Zhan et al., 2023 | 2023 | Potato | 4 |
| 90 | Zhong et al., 2023 | 2023 | Rice | 2 |
| 91 | Zhong et al., 2023 | 2023 | Wheat | 7 |
| 92 | Zhong et al., 2023 | 2023 | Tomato | 7 |
| 92 | Zhong et al., 2023 | 2023 | Larix | 6 |
| 92 | Jalilian et al., 2023 | 2023 | Rice | 6 |
| 92 | Cheng et al., 2023 | 2023 | Rice | 4 |
| 93 | Zhang et al., 2023 | 2023 | Rice | 16 |
| 94 | Biswas et al., 2023 | 2023 | Rice | 4 |
| 95 | Zhou et al., 2023 | 2023 | Rice | 6 |
| 96 | Ahmad et al., 2024 | 2024 | Brassica juncea | 2 |
| 97 | Hui et al., 2024 | 2024 | Cotton | 8 |
| 98 | Yao et al., 2024 | 2024 | Rice | 6 |
| 99 | Vu et al., 2024 | 2024 | Tomato | 3 |
| 100 | Nagy et al., 2025 | 2025 | Maize | 30 |
| 101 | Liu et al., 2025 | 2025 | Rice | 12 |
| 102 | Liu et al., 2025 | 2025 | Tomato | 3 |
| 103 | Choi et al., 2025 | 2025 | Rice | 3 |
| 104 | Contiliani et al., 2025 | 2025 | Rice | 4 |
| 104 | Luo et al., 2025 | 2025 | Tobacco | 6 |
| 105 | Pan et al., 2025 | 2025 | Wheat | 3 |
| 106 | Abdallah et al., 2025 | 2025 | Wheat | 3 |

**Table S2.** Lineage summary per multiplex level.

| Multiplex | Independent lineages (T1) | T2 plants sequenced | T3 plants sequenced | Total plants |
| --- | --- | --- | --- | --- |
| 8 | 36 | 36 | 47 | 83 |
| 16 | 36 | 36 | 30 | 66 |
| 24 | 12 | 12 | 15 | 27 |

**Table S3.** Model comparison for per-site editing efficiency decay. Seven models were fit to plant-level counts of edited sites (Binomial or Beta-Binomial likelihood on number of sites edited / total sites targeted, *n* = 127 plant-by-construct observations). Parameters were estimated by maximum likelihood (Nelder-Mead). 95% confidence intervals are Wald intervals from the inverse observed information (Hessian). Pseudo-R^2^ is McFadden’s relative to an intercept-only null 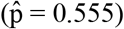. ΔAIC is computed against the best model (M7).

| Mod. | Description | Par. | Estim. | 95% CI | AIC | $\Delta AIC$ | pseudo-$R^2$ |
| --- | --- | --- | --- | --- | --- | --- | --- |
| M1 | $P = p$ | $p$ | 0.730 | fixed | 691.2 | 236.9 | 0.000 |
| M2 | $P = p^n$ | $p$ | 0.555 | [0.530, 0.580] | 13047.7 | 12593.4 | -17.874 |
| M3 | $P = p_0 \cdot n_{base} / n$ | $p_0$ | 0.555 | — | 1007.7 | 553.3 | -0.455 |
| | | $n_{base}$ | 8.000 | fixed | | | |
| M4 | $\text{logit}(P) = a + b \cdot n$ | $a$ | 0.998 | [0.696, 1.301] | 665.6 | 211.3 | 0.043 |
| | | $b$ | -0.049 | [-0.067, -0.031] | | | |
| M5 | $\text{logit}(P) = a + b \cdot \log(n)$ | $a$ | 2.526 | [1.820, 3.232] | 651.5 | 197.2 | 0.063 |
| | | $b$ | -0.856 | [-1.115, -0.598] | | | |
| M6 | M5 + 1% efficiency | $a$ | 2.531 | [1.819, 3.243] | 651.4 | 197.1 | 0.063 |
| | | $b$ | -0.865 | [-1.126, -0.604] | | | |
| **M7** | **M6 + overdispersion** | $a$ | 2.956 | [1.517, 4.396] | **454.3** | **0.0** | **0.351** |
| | | $b$ | -1.030 | [-1.582, -0.478] | | | |
| | | $\rho$ | 0.210 | [0.147, 0.274] | | | |

<sup>1</sup> M3 is weakly identified: $p_0$ and $n_{base}$ trade off along a ridge in the likelihood, so marginal Wald intervals are uninformative (joint CI on the ratio $p_0 \cdot n_{base}$ is narrow).
*Models M1–M3 are analytical nulls; M4–M7 are empirically fitted. Floor $c = 0.01$ in M6 and M7. For M7, $\mu$ is the beta-binomial mean and $\rho$ is the intraclass correlation; individual plant editing probability $p \sim \text{Beta}(\mu(1-\rho)/\rho, (1-\mu)(1-\rho)/\rho)$ . The beta-binomial model improves fit by $\Delta AIC \approx 193$ over M6, reflecting substantial plant-to-plant variation in editing competence that the binomial models cannot absorb.*

**Fig. S1.**
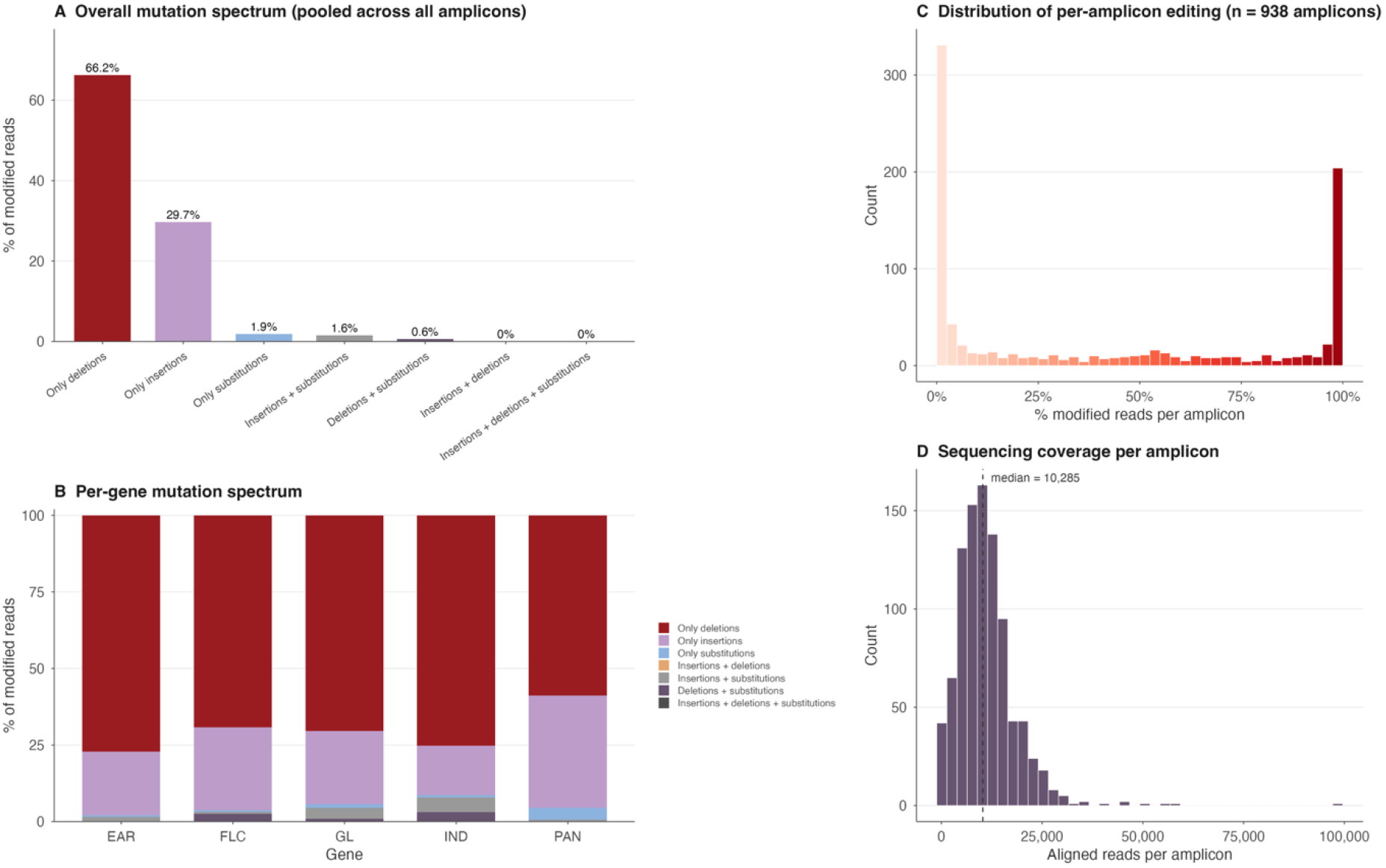
Coverage of amplicon sequencing and mutation spectrum. The figure shows the most common type of mutations across all Amplicons and split by gene. The distribution of reads including a mutation different from the reference sequence and the distribution of coverage per applicants.

**Fig. S2.**
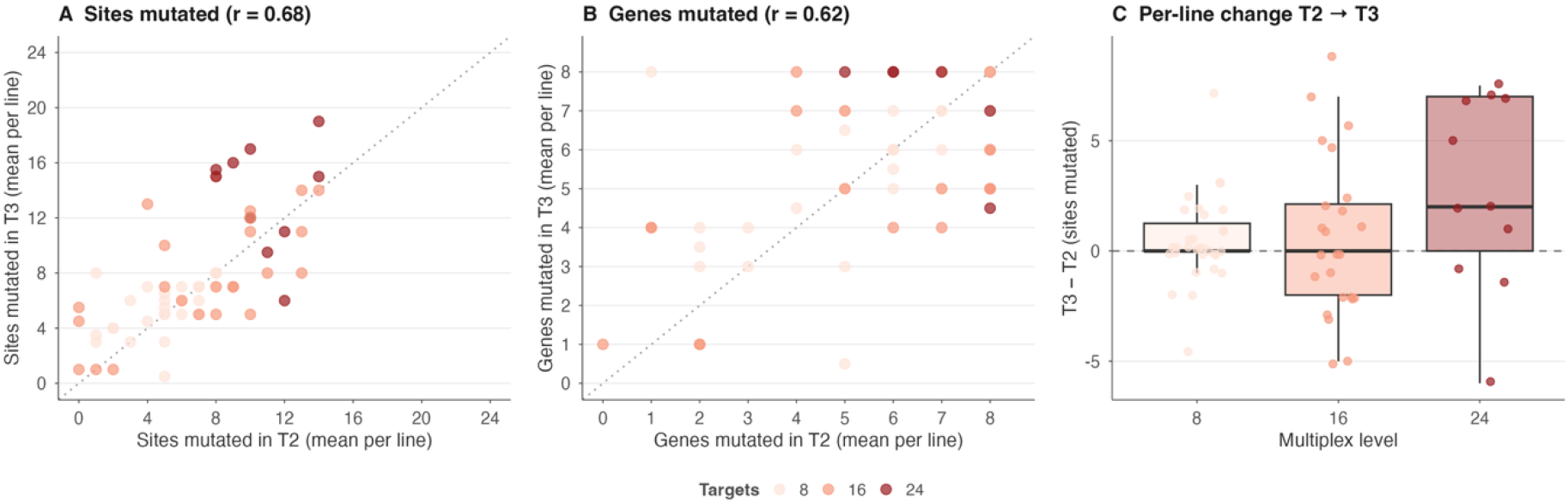
Corresponding number of mutations in T2 and T3 plants. The number of PAM sites mutated or total genes KOs of the same transformed T1 lineage at T2 and T3 sequencing. The difference in sites mutated suggests further Cas9 activity even at T3 that continues increasing the number of mutations.

## References

1. T. Lappalainen, Y. I. Li, S. Ramachandran, A. Gusev, Genetic and molecular architecture of complex traits. Cell 187, 1059–1075 (2024).

2. R. Chen, et al., Multiplex-edited mice recapitulate woolly mammoth hair phenotypes. bioRxiv (2025).

3. L. Impens, et al., Combining multiplex gene editing and doubled haploid technology in maize. New Phytol. 239, 1521–1532 (2023).

4. J.-F. Li, et al., Multiplex and homologous recombination-mediated genome editing in Arabidopsis and Nicotiana benthamiana using guide RNA and Cas9. Nat. Biotechnol. 31, 688–691 (2013).

5. K. Xie, B. Minkenberg, Y. Yang, Boosting CRISPR/Cas9 multiplex editing capability with the endogenous tRNA-processing system. Proc. Natl. Acad. Sci. U. S. A. 112, 3570–3575 (2015).

6. Y. Gao, Y. Zhao, Self-processing of ribozyme-flanked RNAs into guide RNAs in vitro and in vivo for CRISPR-mediated genome editing: Self-processing of ribozyme-flanked RNAs into guide RNAs. J. Integr. Plant Biol. 56, 343–349 (2014).

7. B. P. Kleinstiver, et al., High-fidelity CRISPR-Cas9 nucleases with no detectable genome-wide off-target effects. Nature 529, 490–495 (2016).

8. B. Zetsche, et al., Multiplex gene editing by CRISPR-Cpf1 using a single crRNA array. Nat. Biotechnol. 35, 31–34 (2017).

9. L. C. Laforest, S. S. Nadakuduti, Advances in delivery mechanisms of CRISPR gene-editing reagents in plants. Front. Genome Ed. 4, 830178 (2022).

10. N. M. Gaudelli, et al., Programmable base editing of A•T to G•C in genomic DNA without DNA cleavage. Nature 551, 464–471 (2017).

11. J. G. Doench, et al., Optimized sgRNA design to maximize activity and minimize off-target effects of CRISPR-Cas9. Nat. Biotechnol. 34, 184–191 (2016).

12. J. Stuttmann, et al., Highly efficient multiplex editing: one-shot generation of 8× Nicotiana benthamiana and 12× Arabidopsis mutants. Plant J. 106, 8–22 (2021).

13. R. S. Zou, et al., Massively parallel genomic perturbations with multi-target CRISPR interrogates Cas9 activity and DNA repair at endogenous sites. Nat. Cell Biol. 24, 1433–1444 (2022).

14. E. A. Boyle, Y. I. Li, J. K. Pritchard, An expanded view of complex traits: From polygenic to omnigenic. Cell 169, 1177–1186 (2017).

15. L. Leventhal, M. Ruffley, M. Exposito-Alonso, Planting genomes in the wild: Arabidopsis from genetics history to the ecology and evolutionary genomics era. Annu. Rev. Plant Biol. 76, 605–635 (2025).

16. E. Aliu, et al., Enhancing Agrobacterium-mediated plant transformation efficiency through improved ternary vector systems and auxotrophic strains. Front. Plant Sci. 15, 1429353 (2024).

17. T. Komari, Y. Hiei, Y. Saito, N. Murai, T. Kumashiro, Vectors carrying two separate T-DNAs for co-transformation of higher plants mediated by Agrobacterium tumefaciens and segregation of transformants free from selection markers. The Plant Journal 10, 165–174 (1996).

18. J.-J. Liu, et al., CasX enzymes comprise a distinct family of RNA-guided genome editors. Nature 566, 218–223 (2019).

19. E. E. Ellison, et al., Multiplexed heritable gene editing using RNA viruses and mobile single guide RNAs. Nat. Plants 6, 620–624 (2020).

20. L. Luo, et al., Establishing an immune system conferring DNA and RNA virus resistance in plants using CRISPR/Cas12a multiplex gene editing. Plant Direct 9, e70070 (2025).

21. X. Wu, et al., Rapid adaptation and extinction in synchronized outdoor evolution experiments of Arabidopsis. Science 391, eadz0777 (2026).

22. K. Labun, et al., CHOPCHOP v3: expanding the CRISPR web toolbox beyond genome editing. Nucleic Acids Research 47, W171–W174 (2019).

23. M. G. Murray, W. F. Thompson, Rapid isolation of high molecular weight plant DNA. Nucleic Acids Res. 8, 4321–4325 (1980).

24. J. J. Doyle, J. L. Doyle, A rapid DNA isolation procedure for small quantities of fresh leaf tissue. Phytochemical bulletin (1987).

25. E. Symeonidi, J. Regalado, R. Schwab, D. Weigel, CRISPR-finder: A high throughput and cost-effective method to identify successfully edited Arabidopsis thaliana individuals. Quant. Plant Biol. 2, e1 (2021).

26. K. Clement, et al., CRISPResso2 provides accurate and rapid genome editing sequence analysis. Nat. Biotechnol. 37, 224–226 (2019).

27. N. Bollier, R. Andrade Buono, T. B. Jacobs, M. K. Nowack, Efficient simultaneous mutagenesis of multiple genes in specific plant tissues by multiplex CRISPR. Plant Biotechnol. J. 19, 651–653 (2021).

28. R. Ursache, S. Fujita, V. Dénervaud Tendon, N. Geldner, Combined fluorescent seed selection and multiplex CRISPR/Cas9 assembly for fast generation of multiple Arabidopsis mutants. Plant Methods 17, 111 (2021).

